# *Trans*-species microRNA loci in the parasitic plant *Cuscuta campestris* have a U6-like snRNA promoter

**DOI:** 10.1101/2022.07.06.498962

**Authors:** Collin Hudzik, Sean Maguire, Shengxi Guan, Jeremy Held, Michael J. Axtell

## Abstract

Small regulatory RNAs can move between organisms during pathogenic interactions and regulate gene expression in the recipient. If and how such “*trans*-species” small RNAs are distinguished from normal small RNAs is not known. The parasitic plant *Cuscuta campestris* produces a number of microRNAs that specifically accumulate at the interface between parasite and host, several of which have been demonstrated to have *trans*-species activity. We find that induction of *C. campestris* interface-induced microRNAs was similar regardless of host species, and can be replicated in haustoria stimulated to develop in the complete absence of a host. We also find that the loci encoding *C. campestris* interface-induced microRNAs are distinguished by a common 10 base-pair *cis*-regulatory element. This element is identical to a previously described upstream sequence element used by all plant small nuclear RNA loci. The sequence context of this element strongly suggests U6-like transcription by RNA polymerase III. The element promotes accumulation of interface-induced miRNAs in a heterologous system. This common promoter element distinguishes *C. campestris* interface-induced microRNA loci from other plant small RNAs; other plant small RNA loci are transcribed by polymerases II or IV, and lack any common promoter motifs. Our data suggest that *C. campestris* interface-induced miRNAs are produced in a manner distinct from canonical miRNAs. All confirmed *C. campestris* microRNAs with confirmed *trans*-species activity are interface-induced and possess these features. We speculate that this distinct production may allow these miRNAs to be exported to hosts.

## Introduction

Small regulatory RNAs are 21 to 24 nucleotide RNAs which play an important role as regulators of gene expression. Plants produce at least three distinct classes of small RNAs. MicroRNAs (miRNAs) are usually 21-22 nucleotides in length and derived from single-stranded stem-loop precursors (Chen, 2005; Axtell, 2013). MiRNAs usually regulate specific target mRNAs post-transcriptionally. Canonical plant miRNAs are processed from longer primary transcripts. Canonical plant miRNA precursors in *Arabidopsis thaliana* are transcribed by RNA polymerase II, have variable lengths, and besides a TATA box (Xie et al., 2005) lack any common *cis* regulatory elements. There are also two distinct types of short interfering RNAs (siRNAs), both of which originate from double-stranded RNA precursors. Shorter siRNAs are 21-22 nucleotides long, and mostly function outside of the nucleus in post-transcriptional silencing of endogenous mRNAs, aberrant RNAs, and viral RNA (Fei et al., 2013). Longer siRNAs are mostly 24 nucleotides long and usually function in the nucleus to direct *de novo* DNA methylation to transposons and other non-genic sequences (Matzke and Mosher, 2014). *Trans*-species small RNAs, defined as small RNAs that naturally move from one organism to another, play important roles in plant-pathogen interactions by targeting mRNAs in the recipient organism (Huang et al., 2019; Hudzik et al., 2020). Most experimentally verified cases involve miRNAs (Zhang et al., 2016; Shahid et al., 2018) or 21-22 nucleotide siRNAs (Weiberg et al., 2013; Cai et al., 2018; Hou et al., 2019).

*Cuscuta campestris* is an obligate parasitic plant which uses a specialized organ called the haustorium to pierce the host stem and fuse with the vasculature to feed (Heide-Jørgensen, 2008). The haustorium facilitates bidirectional movement of photosynthates and macromolecules between the host and parasite (Birschwilks et al., 2006; Roney et al., 2007; David-Schwartz et al., 2008; Kim and Westwood, 2015). Haustorium organogenesis is divided into the adhesive, intrusive, and conductive phases (Shimizu and Aoki, 2019). During the adhesive phase, cortical cells from the parasite stem begin to differentiate and divide to form the endophyte primordia. Digitate cells from the endophyte primordia eventually extend through and invade the host stem, marking the transition into the intrusive phase. Digitate cells, now referred to as searching hyphae, continue to extend and move in-between cortical cells to identify the host vasculature (Vaughn, 2003). Searching hyphae will differentiate into xylic or phloic hyphae once in contact with the respective vasculature. Haustoria have fully matured and reached the conductive phase once differentiated hyphae begin to actively receive nutrients from the host (Vaughn, 2006).

*C. campestris* lacks several miRNA families that are otherwise common in closely related species (Zangishei et al., 2022). Other *C. campestris MIRNA* loci may have been acquired through horizontal gene transfer (Yang et al., 2019; Zangishei et al., 2022). These patterns of loss and gain mirror those observed for *Cuscuta* protein-coding genes (Sun et al., 2018; Vogel et al., 2018). *C. campestris* produces a set of miRNAs specifically at the interface between the host and parasite (Shahid et al., 2018; Johnson et al., 2019), some of which can move long distances in the host plant (Subhankar et al., 2021). Several interface-induced miRNAs from *C. campestris* have been experimentally demonstrated to target and regulate host mRNAs, and thus act in a *trans*-species manner. Host mRNAs involved in biotic defense, hormone signaling, and vascular development are targeted (Shahid et al., 2018; Johnson et al., 2019). A subset of the interface-induced, *trans*-species miRNAs are 22 nucleotides long, which allows for production host-derived secondary siRNAs to accumulate during parasitism (Chen et al., 2010; Cuperus et al., 2010). Analysis of interface-induced, *trans*-species miRNA diversity within and between *Cucuta* species provides clear evidence that several miRNA-target relationships have been under selection (Johnson et al., 2019). When *C. campestris* is grown on a host plants with a non-functional copy of certain targets parasite biomass increases significantly (Shahid et al., 2018). Thus we hypothesize that *C. campestris* uses interface-induced, *trans*-species miRNAs to manipulate host gene expression to increase parasite fitness. We use the term “interface-induced miRNAs” to refer to all such differentially expressed miRNAs because not all of them have been directly proven to target host mRNAs.

In this study we examine the regulation of *C. campestris* interface-induced miRNAs with respect to haustorial development and host identity. We also analyze *cis*-regulatory features found at the loci encoding *C. campestris* interface-induced miRNAs.

## Results

### Improved annotation of interface-induced *MIRNA* loci from *C. campestris*

A total of 43 *C. campestris* interface-induced *MIRNA* loci, representing 36 distinct families, were previously identified based on a single small RNA-seq experiment with two biological replicates per tissue type (Shahid et al., 2018). Subsequently several more small RNA-seq datasets from *C. campestris* infestation of *A. thaliana* have been produced ((Johnson et al., 2019); Supplemental Table 1). A *C. campestris* draft genome assembly with higher contiguity relative to the one used by (Shahid et al., 2018) has also been described (Vogel et al., 2018). The additional small RNA-seq data were used in conjunction with the improved genome assembly to identify interface-induced *MIRNA* loci. A total of 156 *C. campestris* interface-induced *MIRNA* (“ccm-IIM”) loci, representing 103 families, were annotated (Supplemental Datasets 1-3). None of these families have been annotated in other plant species (per miRBase version 22). These improved annotations more than triple the known number of *C. campestris* interface-induced *MIRNA* loci, and include all miRNAs previously shown to target host mRNAs. A total of 21 out of the 103 families have one or more experimental lines of evidence supporting *trans*-species activity against host mRNAs (Shahid et al., 2018; Johnson et al., 2019; Figure 1; Supplemental Table 2). All data are searchable with genome browsers and integrated small RNA visualization tools at https://plantsmallrnagenes.science.psu.edu/Studies/Hudzik.

**Figure 1:**
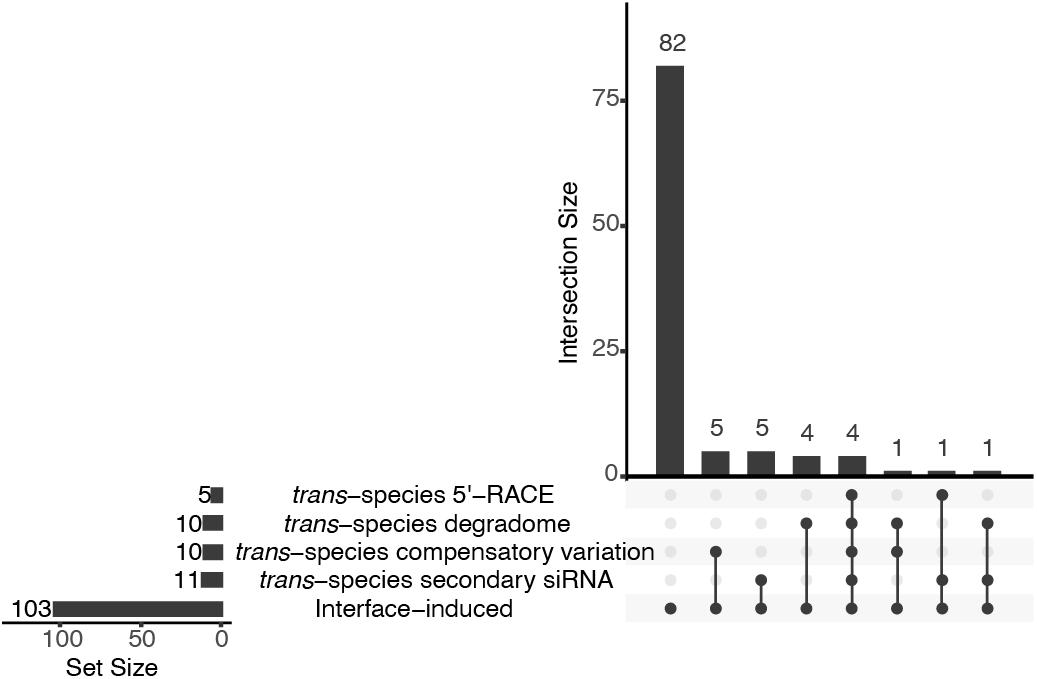
Evidence for *trans*-species targeting of host mRNAs by *Cuscuta campestris* interface-induced miRNA families. See Supplemental Table 2 for details.

### *C. campestris* interface-induced miRNAs accumulate prior to host penetration

Whether using *A. thaliana* inflorescence stems or *Solanum lycopersicum* hypocotyls as a host, haustorium organogenesis progressed at the same rate (Supplemental Figures 1–2). Haustoria were in the adhesive phase during days 0-2 of the time-course when the formation of the endophyte primordia was observed (Figures 2A, 2D). The intrusive phase, discerned by searching hyphae invading the host cortex, spanned days 3-5 (Figures 2B, 2E). A xylem bridge connecting host and parasite vasculature via the haustorium was not detectable until day 6 (Figures 2C, 2F); this marks the start of the conductive phase.

**Figure 2:**
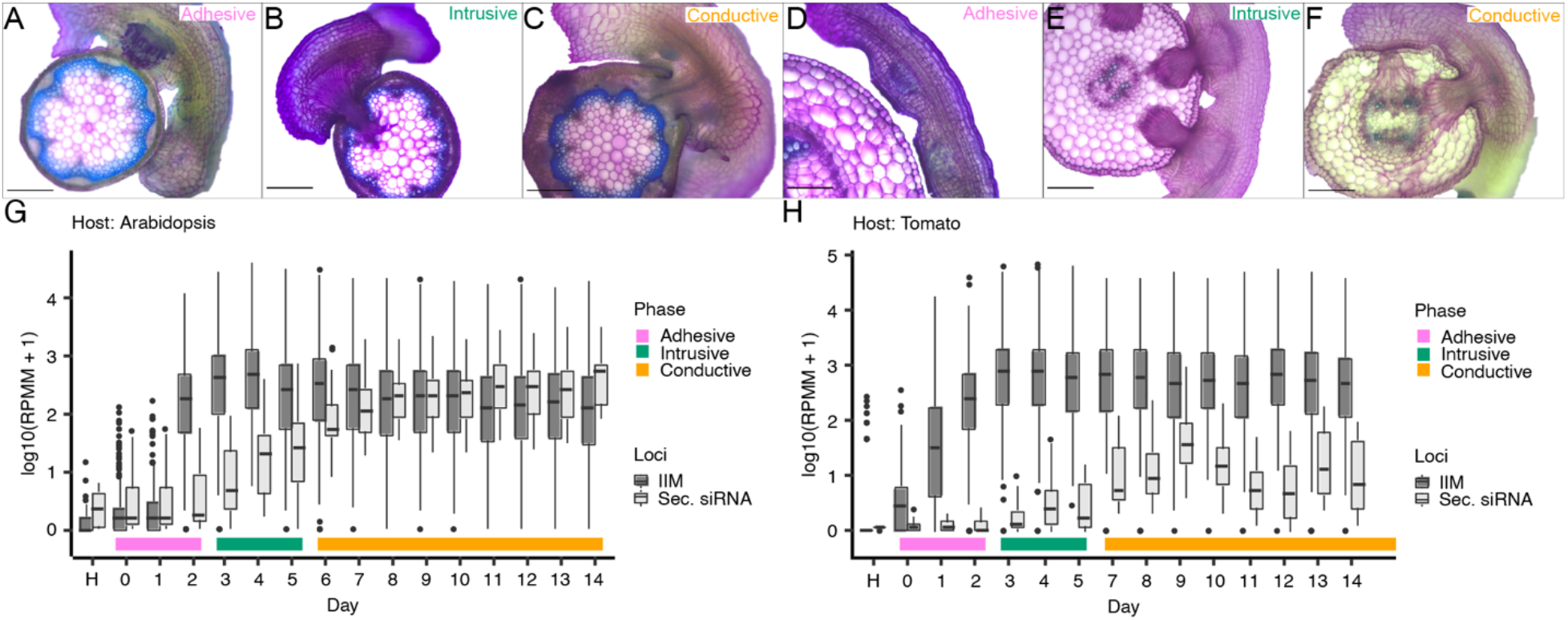
Interface-induced miRNAs are detectable during the adhesive phase. Histological sections of haustoria growing on *A. thaliana* (A-C) and *S. lycopersicum* (D-F). Top right text in each image (A-F) denotes the stage of haustorium organogenesis, the color correlates with the phases labeled in G and H. (Scale bars: 200μm) (G, H) Detection of interface-induced miRNAs (IIM) and secondary siRNAs (Sec. siRNA) using *A. thaliana* or *S. lycopersicum* as hosts. Boxplots show medians (central line), 25^th^-75^th^ percentiles (box boundaries), up to 1.5x the interquartile range (whiskers), and outliers (dots). RPMM: Reads per million mapped. Histological sections from the full time-course, including the images shown in A-F, are shown in Supplemental Figures 1–2.

sRNA sequencing was performed across a 15-day time-series (days 0-14) of *C. campestris*-host interfaces for both *A. thaliana* and *S. lycopersicum* hosts. Interface-induced miRNAs began accumulating in the adhesive phase, and reached maximum levels by the start of the intrusive phase (Figure 2G, 2H). Interface-induced miRNAs can stimulate the generation of secondary siRNAs following an initial cleavage event on a host transcript (Shahid et al., 2018; Johnson et al., 2019). Appearance of secondary siRNAs must therefore occur after miRNAs interact with host mRNAs. Accumulation of secondary siRNAs did not begin until the intrusive phase, and did not reach maximum levels until the conductive phase (Figure 2G, 2H). These data show that interface-induced miRNAs appear early in haustorial development, before any penetration of host tissue has occurred.

### *C. campestris* interface-induced miRNA induction is not host-dependent

*C. campestris* has a broad host range. One possible method of adapting to diverse hosts could be to adjust interface-induced miRNA accumulation depending on the host species. This hypothesis was tested by plotting the median conductive phase small RNA accumulation levels of each interface-induced *MIRNA* locus from tomato and *A. thaliana* hosts. The resulting plot showed a linear correlation (Figure 3A), where each *MIRNA* locus accumulated miRNAs to similar levels with either species as a host. This argues against the hypothesis that miRNA induction varies depending on the host species.

**Figure 3:**
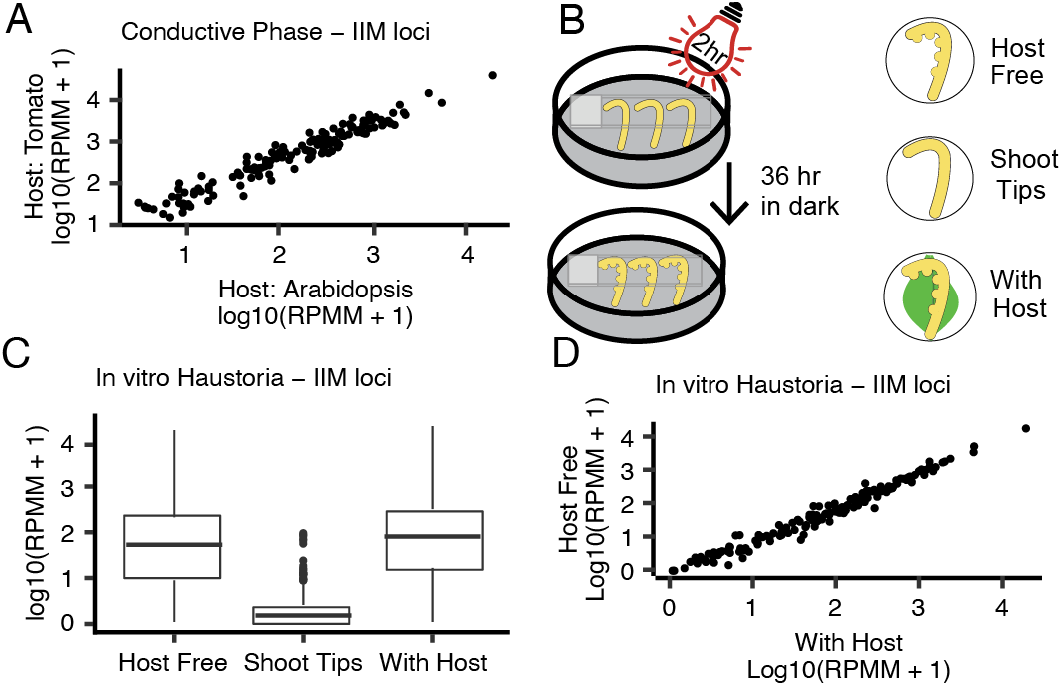
Interface-induced miRNA production is independent of a host. (A) Comparing interface-induced miRNA population and abundance between hosts. Each dot represents the median value of conductive phase samples for an interface-induced miRNA from the sRNA-seq time-course. (B) Graphical overview of *in vitro* haustoria experiment. (C) Interface-induced miRNA accumulation from *in vitro* haustoria. Boxplots show medians (central line), 25^th^-75^th^ percentiles (box boundaries), up to 1.5x the interquartile range (whiskers), and outliers (dots). (D) Comparing interface-induced accumulation between the Host Free and With Host *in vitro* haustoria. Each dot represents the median value across all replicates for a single interface-induced microRNA from sRNA-seq of *in vitro* haustoria. IIM: Interface-induced microRNA; RPMM: Reads per million mapped.

Detached shoot tips of *C. campestris* can be stimulated to differentiate haustoria by specific light and tactile stimuli (Figure 3B; (Kaga et al., 2020)). These *in vitro* haustoria can be produced either with or without adjacent host tissue, such as detached host leaves (Figure 3B). *In vitro* haustoria produced with or without adjacent *A. thaliana* leaves were processed in triplicate for small RNA sequencing to test interface-induced miRNA accumulation. *C. campestris* shoot tips (also in triplicate) were used as a negative control. Interface-induced miRNAs were abundant in *in vitro* haustoria regardless of the presence or absence of detached host leaves (Figure 3C). Median accumulation levels of individual miRNAs were highly correlated between *in vitro* haustoria produced with or without adjacent host tissue (Figure 3D). These observations demonstrate that *C. campestris* interface-induced miRNA production is independent of any host. Instead, their sudden appearance during early haustorial development is part of an inherent developmental program.

### *C. campestris* interface-induced *MIRNA* loci share a common upstream sequence element known to control snRNA transcription in plants

Genomic DNA sequences upstream of the interface-induced loci were analyzed with MEME (Bailey et al., 2009) to search for over-represented sequence motifs. Transcription start sites for these *MIRNA* primary transcripts were initially unknown, so the ends of the upstream regions were defined by the 5’ basal DCL cut site on the *MIRNA* hairpins (Figure 4A). A highly overrepresented ten nucleotide motif was discovered (Figure 4B). This motif, termed the Upstream Sequence Element (USE), was found adjacent to most (145/156) of the interface-induced *MIRNA* loci (Figure 4C). As a control, a set of 37 “canonical” *MIRNA* loci from *C. campestris* were annotated (Supplemental Dataset 4). These canonical loci encode miRNAs universal among dicots such as miR156, miR164, and miR166. None of the canonical *MIRNA* loci had the USE in their upstream regions (Figure 4C). The motif has a strong tendency to be located ~100 base pairs upstream of the 5’ basal DCL cut sites of the interface-induced *MIRNA*s (Figure 4D). A previous study produced nanoPARE data from *C. campestris* / *A. thaliana* haustorial interfaces (Johnson et al., 2019); nanoPARE is an RNA-seq method that captures 5’ ends of RNAs. These data were analyzed to find the 5’ ends of *MIRNA* primary transcripts. There was a clear peak of nanoPARE data around 40 base-pairs upstream of the 5’ basal DCL cut site of the interface-induced *MIRNA*s (Figure 4E). No such peak was apparent when analyzing the canonical *MIRNA* loci (Figure 4E). This analysis suggests that the transcriptional start sites of interface-induced *MIRNA* primary transcripts about 40 bps upstream of the 5’-basal DCL cut site, and thus lie downstream of the USE.

**Figure 4:**
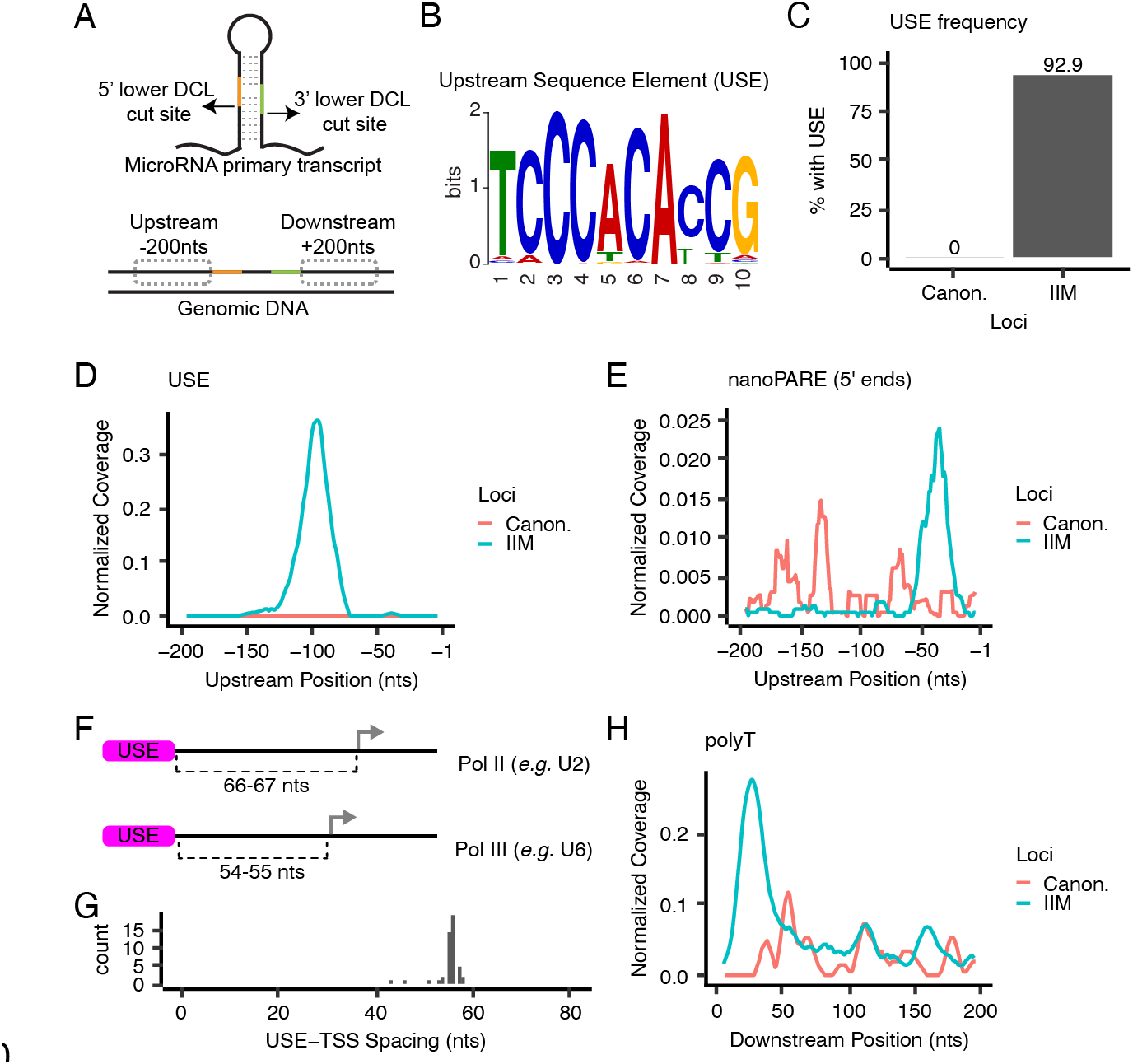
Interface-induced *MIRNA* loci share a *cis*-regulatory element with snRNAs. (A) Schematic showing a *MIRNA* primary transcript (top), and the corresponding genomic locus (bottom). Upstream and downstream regions were anchored by the lower Dicer-Like (DCL) cut sites of the primary transcript. (B) MEME sequence logo of the USE found upstream of interface-induced *MIRNA* loci. (C) Presence of the USE at interface-induced *MIRNA* loci and canonical *MIRNA* loci in *C. campestris*. (D) Metaplot of USE coverage as a function of upstream position. (E) Metaplot of normalized nanoPARE 5’ end coverage. (F) Distances between USE and transcriptional start sites (arrows) for known USE-dependent Pol II and III type promoters. (G) Frequency distribution of USE-TSS distances for interface-induced miRNAs. (H) Metaplot of polyT coverage (defined as six or more consecutive T residues on the coding strand) in downstream regions. Canon.: Canonical microRNA loci; IIM: Interface-induced microRNA loci; TSS: Transcriptional start site.

### *C. campestris* interface-induced *MIRNA* loci have features consistent with RNA Polymerase III transcription

The USE found at *C. campestris* interface-induced *MIRNA* loci is identical to a motif known to drive U-snRNA transcription in plants (Vankan and Filipowicz, 1989). U-snRNAs are short non-coding RNAs that, along with associated proteins, form the spliceosome. The spliceosome functions to catalyze intron removal from Pol II-transcribed RNAs during nuclear RNA maturation. The USE is a unique *cis*-regulatory element because it can drive either Pol II or Pol III transcription of U-snRNAs (Waibel and Filipowicz, 1990b). When the USE is 66-67 base-pairs upstream from the transcriptional start site (TSS) it drives Pol II transcription (Figure 4F). When the USE is 54-55 base-pairs upstream of the TSS (one helical turn of DNA shorter), it instead drives Pol III transcription (Figure 4F). *A. thaliana* U2, U4, and U5 snRNA genes contain the longer spacing and are transcribed by Pol II, while *A. thaliana* U6 snRNA genes have the shorter spacing and are transcribed by Pol III (Waibel and Filipowicz, 1990b; Waibel and Filipowicz, 1990a). The TSSs of 48 of the *C. campestris* interface-induced MIRNA primary transcripts were inferred from nanoPARE data (Figure 4E); the rest could not be inferred due to low/no coverage in the nanoPARE data. The USE-TSS distance at these loci had a sharp peak at about 55 base-pairs (Figure 4G), exactly the spacing expected for USE-dependent Pol III transcription. Pol III termination is triggered by runs of T residues on the non-template strand; four Ts is a minimal signal for termination, and six or more Ts are maximally effective (Gao et al., 2018). Runs of six or more T’s were common immediately downstream of interface-induced *MIRNA* loci (Figure 4H). The upstream similarity to U6 snRNA promoters and the prevalence of downstream poly-T stretches suggests that *C. campestris* interface-induced *MIRNA* loci could be transcribed by Pol III.

### Accumulation of interface-induced microRNAs in a heterologous system depends on the USE

*Agrobacterium tumefacians* T-DNA vectors containing *C. campestris* interface-induced *MIRNA* loci were constructed (Figure 5A). The selected miRNAs have both been previously shown to have *trans*-species silencing activity on host mRNAs: miR12463 targets *BIK1* and miR12497 targets *TIR1* and related *AFB* mRNAs (Supplemental Table 2; Shahid et al., 2018, Johnson et al. 2019). The T-DNA did not contain any selectable markers to avoid recruitment of RNA polymerases; transcription of these loci depends solely on any *cis* elements present in the *MIRNA* loci. For each *MIRNA* tested a wild-type and USE-scrambled version were prepared. These T-DNA were introduced to *Nicotiana benthamiana* leaf mesophyll cells by Agroinfiltration. Mature miRNA levels were quantified by qRT-PCR. Mature miRNAs accumulated when wild-type loci were introduced (Figures 5B-C). When the USE element was scrambled, mature miRNA accumulation was dramatically lower (Figures 5B-C). This supports the hypothesis that the USE is a *cis*-acting factor that promotes miRNA accumulation. This result also demonstrates the *C. campestris* interface-induced *MIRNA* precursors can be correctly processed by the RNAi machinery of a heterologous host.

**Figure 5:**
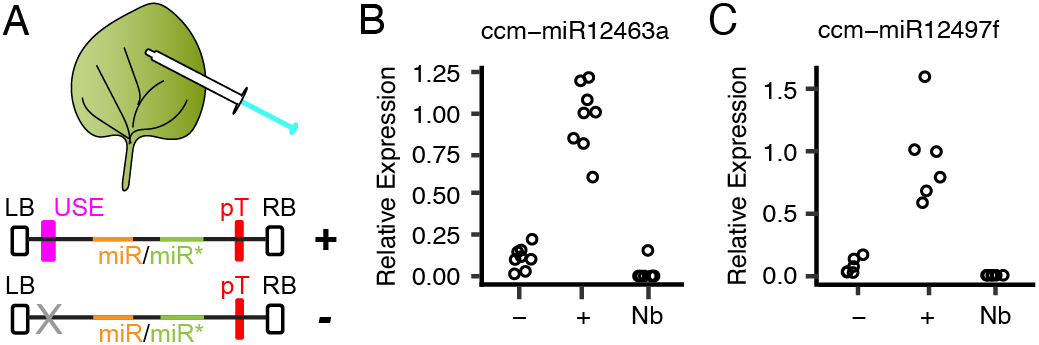
The USE is required for accumulation of interface-induced miRNAs during transient expression in *N. benthamiana*. (A) Graphical representation of constructs used during transient expression. LB: left border; USE: upstream sequence element; pT: polyT stretch; RB: right border. (B-C) Relative expression of miR12463a or miR12497f with scrambled USE sequence (−), wild-type USE sequence (+), and *N. benthamiana* leaf not infiltrated (Nb). Expression was normalized to miR159. Each dot represents a biological replicate (RNAs from a distinct infiltrated leaf).

## Discussion

We observed that *C. campestris* interface-induced miRNAs begin accumulating in the adhesive phase, even before the parasite has penetrated the host stem. This suggests that interface-induced miRNAs are primed for immediate use by developing haustoria. Accumulation of interface-induced miRNAs is fully independent of the host identity and does not even require any host. This indicates that their accumulation is a programmed aspect of haustorial development. *C. campestris* is a generalist parasite that feeds on a wide range of host plants. This “shotgun” strategy of miRNA accumulation could be well-suited for a generalist that feeds on diverse hosts. This host-insensitive, shotgun strategy is also consistent with earlier observations of *Cuscuta* miRNA polymorphisms that compensate for most possible variations of target sites within host mRNAs (Johnson et al., 2019).

We also found a snRNA-like USE adjacent to nearly all interface-induced *MIRNA* loci. The USE is not found at any canonical miRNA loci in *C. campestris* and has not been reported at *MIRNA* loci in any other plant. The distance between the USE and transcriptional start sites of interface-induced miRNAs is consistent with U6-like Pol III transcription, as is the presence of potential Pol III terminators immediately downstream of most loci. The USE drives accumulation of miRNAs in a heterologous system, directly demonstrating its function as an activating *cis*-regulatory element. We hypothesize that the *C. campestris* interface-induced miRNA precursors are transcribed by RNA Pol III in a USE-dependent manner. All confirmed *trans*-species miRNAs from *C. campestris* are also interface-induced and contain the USE. Only 20% of the interface-induced miRNA families have thus far been proven to target host mRNAs (Figure 1). However, many other targeting events may be undetected. Detection relies on stable sliced remnants of host mRNAs, or on appearance of secondary siRNAs. Some miRNA-target interactions do not cause either outcome. Given their coordinated expression and common *cis*-regulatory motifs, we suspect that many more of the interface-induced miRNAs do in fact target host mRNAs during *C. campestris* parasitism.

We speculate that the use of Pol III, instead of Pol II, may somehow mark *trans*-species miRNAs for specialized processing and eventual export from the parasite into the host. The *trans*-species miRNAs avoid “self-targeting” of *C. campestris* mRNAs (Shahid et al., 2018; Johnson et al., 2019); this is consistent with the hypothesis that they are made for “export only” and not assembled onto *C. campestris* AGO proteins. The accumulation of mature miRNAs in *in vitro* haustoria with no host tissue present directly demonstrates that dicing of the *trans*-species miRNA precursors occurs within *C. campestris*. Therefore, the exported molecule is either mature miRNA or the miRNA/miRNA* duplex.

Plant U6 promoters are workhorses for constitutive expression of short non-coding RNAs. For instance, the U6-26 promoter from *A. thaliana* is widely used to drive Pol III-dependent sgRNA expression in CRISPR experiments (Tsutsui and Higashiyama, 2017; Lowder et al., 2018). U6-26 has the same USE as the *C. campestris* interface-induced miRNAs (Waibel and Filipowicz, 1990a). Indeed, the U6 snRNA is often used as a loading control for miRNA blots, including in the first report of interface-induced miRNAs (Shahid et al., 2018). How does the USE, which in all previously known contexts drives constitutive transcription, result in miRNAs with such a tissue-specific accumulation pattern? One hypothesis is constitutive transcription but tissue-specific dicing of the precursors. However, RNA blots and RNA-seq experiments have yet to detect any strong precursor accumulation in non-haustorial tissues. Resolving the mystery of highly specific accumulation from a promoter that is expected to be constitutive will be an important goal for future research.

In summary, we find that the *C. campestris* interface-induced *MIRNA* loci are distinguished from canonical *MIRNA* loci by a distinct set of genomic sequence features. This implies that these miRNAs are fundamentally distinct from normal miRNAs and siRNAs. The common promoter element suggests a possible avenue for disruption of *trans*-species miRNA activity: targeting the factors that bind to the USE should reduce production of *trans*-species miRNAs.

## Methods

### Annotation of *Cuscuta campestris* interface-induced *MIRNA* loci

Annotation of *MIRNA* loci occurred in two steps (early and late) based on the availability of small RNA-seq data during the course of the study. In the early step, previously published small RNA-seq datasets from the interface or parasite stem of *C. campestris* attached to *A. thaliana* were obtained as fastq files from the Sequence Read Archive (SRA; Supplemental Table 1). Trailing 3’-sequencing adapters were discovered and trimmed using gsat (https://github.com/MikeAxtell/gsat). Trimmed small RNA datasets were then aligned to the *A. thaliana* genome (version TAIR10, including plastid and mitochondrial genomes) using bowtie (Langmead et al., 2009) demanding exact matches and directing unmapped reads to a new fastq file (non-default settings ‘-v 0 --un’). Small RNA reads that aligned were discarded; this eliminated most small RNAs that could have been produced by the host genome. The remaining reads were analyzed together in a single ShortStack run (version 3.8.5; default settings) (Johnson et al., 2016) using the *C. campestris* genome assembly version 0.32 (Vogel et al., 2018) as the reference genome. This resulted in *de novo* annotation of thousands of discrete small RNA-producing loci. Read counts of aligned small RNAs from each locus in each library were retrieved (from the ‘Counts.txt’ file produced by ShortStack) and used as input for differential expression analysis using DESeq2 (Love et al., 2014). Differential expression analysis of interface vs. parasite stem was independently performed for each of the three studies included in the first step (Shahid et al. expt.1, Shahid et al. expt. 2, and Johnson et al.; Supplemental Table 1). DESeq2 analysis used ‘lfcShrink’ using the ‘normal’ shrinkage estimator; loci up-regulated in interface relative to parasite stem with an FDR-adjusted p-value of < 0.05 were called differentially expressed. The union of all three sets of differentially expressed loci from each study was then computed and retained. From these, loci where a) greater than 80% of aligned reads were between 20-24 nucleotides in length and b) the single most abundant aligned RNA size was 20, 21, 22, or 23 nucleotides long were retained. The retained loci were then individually examined using multiple visualization tools: Integrative Genomics Viewer (Robinson et al., 2011), strucVis (https://github.com/MikeAxtell/strucVis), and sRNA_Viewer (https://github.com/MikeAxtell/sRNA_Viewer). Based on the manual examinations, loci that did not conform to all of the guidelines for confident plant *MIRNA* locus annotation (Axtell and Meyers, 2018) were discarded. The late phase of *MIRNA* annotation took advantage of the much greater amount of sRNA-seq data generated during the time-course analyses of *C. campestris* on both *A. thaliana* and *S. lycopersicum* hosts (Supplemental Table 1). The same analysis approach was used to identify interface-induced *MIRNA* loci using days 9 and 10 as ‘Interface’ samples and day 0 and “host only” as negative control samples. Loci identified in the late phase were then merged with those obtained in the early phase. All together 156 *C. campestris* interface-induced *MIRNA* loci were found (Supplemental Datasets 1-3). These loci and the supporting data can be interactively browsed at https://plantsmallrnagenes.science.psu.edu/Studies/Hudzik.

The mature 5p-miRNA and 3p-miRNA from each locus was queried against miRBase (version 22) mature miRNAs using the SSEARCH function (hosted on the miRBase website), restricting hits to Viridiplantae. The mature 5p-miRNA and 3p-miRNA from each loci were also queried against the “superfamilies” described by (Johnson et al., 2019) using ggsearch36 (from the FASTA package), restricting the search to forward strand hits only; alignments were kept that had less than 3 mismatches. In order to group the loci into families, The mature 5p-miRNA and 3p-miRNA from each loci were queried against each other in an all vs. all search using ggsearch36, restricting the search to forward strand hits only. Families were defined by no more than 7 mismatches between a given 5p-miRNA and 3p-miRNA. Based on this analysis the 156 loci were grouped into 103 distinct families^1^.

### Plant growth conditions

Host plants grown for time-course experiments, *A. thaliana* (Col-0) and *S. lycopersicum* (IL-8-1-1), were grown at 20-22°C and subjected to 16h photoperiods under cool-white fluorescent tube lighting. *C. campestris* seeds were scarified with sulfuric acid for 1 hour, followed by 5-6 washes with water. Scarified seeds were placed in glass petri dishes with moist paper towels and placed in a growth chamber at 28°C with 16h photoperiods under fluorescent tube lighting for 3 days. Emerged seedlings were then transplanted to the base of host plants and supplemented with far-red LED lighting throughout the time-course to allow for attachment.

### Time-course of *C. campestris* haustoria development

Host plants were grown as described above until inflorescence stems of *A. thaliana* hypocotyls of *S. lycopersicum* were able to serve as hosts for *C. campestris. C. campestris* seedlings were scarified and germinated as described above and were placed at the base of inflorescence stems and hypocotyls under a combination of cool-white fluorescent tube lighting and far-red LED lights. Day 0 of the time-course was defined as the day that *C. campestris* had successfully wrapped around a host. Samples were subsequently collected every twenty-four hours. Each day in the time course included three biological replicates consisting of four interfaces. Samples which were used to create sRNA sequencing libraries were flash frozen in liquid nitrogen and stored at −80°C until total RNA extraction was performed using a BioSpec Mini-Beadbeater and Tri-Reagent (Sigma) following the manufacturer’s protocol.

Samples which were subjected to vibratome sectioning were collected and sectioned within the same day. A .stl file created by (Atkinson and Wells, 2017) containing instructions for 3D printing a mold to embed samples in agarose for vibratome sectioning was printed. Interfaces were placed in the mold and embedded in freshly prepared 5% agarose. Once solidified, excess tissue and agarose were trimmed, and the samples were mounted on a metal block using superglue. Mounted samples were secured with a vice grip in the Vibratome 1000 Plus Sectioning System bath. Samples were submerged in ice-cold nanopure water and sectioned with a thickness of 200μm using Personna 3-Facet Double Edge AccuTec Blades at a blade angle of 15°. Sections were stained with freshly prepared 0.002% toluidine blue on a shaker for ten minutes in 3×4 cell culture plates and rinsed 3-4 times with nanopure water. Sections were observed on a Zeiss AXIO Scope A1 trinocular optical microscope using the 10x objective and imaged with a Jenoptik ProgRes C14 Plus microscope CCD. Image processing to remove excess stained agarose from the background was identical for all images. Scale bar of 200μm was added to all images using ImageJ.

### *In vitro* haustoria

Stimulation of haustorial development in the absence of host tissues was similar to previously described methods (Kaga et al., 2020; Jhu et al., 2021). *C. campestris* was cultivated on beets in a greenhouse. Shoot tips, approximately 6-8 cm in length, were cut and placed on a 3% agarose plate. Unstimulated shoot tips were flash-frozen at this point for later RNA extraction. Shoot tips “with host” included fresh cut rosette leaves from wild-type (Col-0) *A. thaliana* plants pressed on the tips. Shoot tips were stimulated to form haustoria by weighting with ~8 glass cover slips, exposure to a far-red LED bulb for two hours, followed by four days in complete darkness. The resulting haustoria were collected, flash-frozen, and used for RNA extraction. Total RNA was extracted using a bead-beater and Tri-Reagent (Sigma) per the manufacturer’s instructions. Three samples per condition, each from a different plate, were collected and extracted.

### Small RNA sequencing and analysis

All small RNA-seq libraries were constructed using 500ng of total RNA using the splint-ligation method described by (Maguire et al., 2020). Libraries were sequenced on an Illumina NextSeq 550 (time-course experiments) or NextSeq 2000 (*in vitro* haustoria experiments). Time-course experiment: After de-multiplexing, FASTQ files were trimmed using cutadapt (Martin, 2011) with settings -a AGATCGGAAGAGCACACGTCTGAAC -m 15 --discard-untrimmed. Two concatenated genomes were prepared: *A. thaliana* TAIR10 and *C. campestris* v0.32, and *S. lycopersicum* release 5 and *C. campestris* v0.32. Trimmed reads were aligned to the appropriate concatenated genome using ShortStack (Johnson et al., 2016) version 3.8.5 with setting --align_only. The resulting BAM-formatted alignments were split based on reference genome, to create two BAM files per time-course: One for *C. campestris*, the other for the host genome. These are the alignments hosted at https://plantsmallrnagenes.science.psu.edu/Studies/Hudzik. The *C. campestris* alignments were analyzed using ShortStack version 3.8.5 with a ‘locifile’ containing the coordinates of all 156 interface-induced *MIRNA* loci (Supplemental Dataset 1). The host-genome alignments were similarly analyzed with a ‘locifile’ containing locations of known secondary siRNA loci triggered by interface-induced miRNAs (Supplemental Table 3). The resulting count matrices, showing alignments by sample, were combined and processed to yield normalized values in units of read per million mapped. Analysis of small RNA-seq from *in vitro* haustoria was identical except that alignments used only the *C. campestris* genome and no secondary siRNA loci were analyzed.

### Analysis of flanking DNA

Upstream and downstream DNA sequences (200 base-pairs each) flanking the 156 interface-induced *MIRNA* loci were retrieved from the version 0.32 *C. campestris* genome assembly (Vogel et al., 2018). Upstream regions were defined to end at the 5’ basal DCL cut site; downstream regions were defined to begin at the 3’ basal DCL cut site (Figure 4A). Upstream sequences were analyzed with MEME suite (version 5.4.1) (Bailey et al., 2009). Motif discovery with meme used settings ‘-dna -mod zoops -nmotifs 1 -w 10 -objfun classic-revcomp-markov_order 0’. The top-scoring motif was then used as a query of the upstream sequences with the MEME suite tool fimo using settings ‘--qv-thresh --thresh 0.05’; this captured all occurrences of the motif with a false-discovery rate of <= 0.05. In a few cases where the motif was found more than once on the same upstream sequence, only the top-scoring instance was retained. The motif consensus sequence is TCCCACA[CT]CG (Figure 4B). The fimo analysis was also run using the upstream regions of conserved, canonical *MIRNA* loci from *C. campestris*; no motif instances were found with an FDR <= 0.05. Motif locations were converted to gff3 format, and then converted to modified gff file with metaplot coordinates based on distance from the 5’ basal DCL sites. The subcommand ‘genomecov’ from the bedtools suite (Quinlan and Hall, 2010) was then used with option -d to calculate position-specific depths. These values were then normalized by dividing by the number of loci in each set (interface-induced *MIRNA* loci or conserved canonical *MIRNA* loci). Moving averages with a window size of 10 were then plotted (Figure 4D). Downstream regions were searched for stretches of six or more T nucleotides on the *MIRNA* coding strand; locations were converted to gff3 format and processed as described above to create metaplots (Figure 4H).

### NanoPARE analysis and transcriptional start site inference

Previously described nanoPARE data (Johnson et al., 2019) from *C. campestris* haustorial interfaces with *A. thaliana* stems were retrieved in FASTQ format from the Sequence Read Archive (SRA): Accessions SRR9216111, SRR9216113, and SRR9216114. Contaminating adapter sequences were removed using cutadapt (Martin, 2011) with settings -a CCGAGCCCACGAGACTAAGGCGAATCTCGTATGCCGTCTTCTGCTTG -a CCGAGCCCACGAGACCGTACTAGATCTCGTATGCCGTCTTCTGCTTG -a CCGAGCCCACGAGACAGGCAGAAATCTCGTATGCCGTCTTCTGCTTG -a CCGAGCCCACGAGACGGACTCCTATCTCGTATGCCGTCTTCTGCTTG -a CCGAGCCCACGAGACTAGGCATGATCTCGTATGCCGTCTTCTGCTTG --trim-n -m 20.

Trimmed reads were aligned to the version 0.32 *C. campestris* genome assembly (Vogel et al., 2018) using STAR (Dobin et al., 2013) with settings --outFilterMismatchNmax 999 --outFilterMismatchNoverReadLmax 0.07 --outWigType wiggle read1_5p --outWigNorm None. The two resulting .wiggle formatted files contained the depths of RNA 5’ ends genome-wide; these files were converted to bigwig format using wigToBigWig (Kent et al., 2010). The bigwig files were then queried to retrieve data from the 200 base-pair upstream regions of each of the interface-induced *MIRNA* loci and each of the conserved canonical *MIRNA* loci using bigWigToBedGraph (Kent et al., 2010). The read depths for each locus were scaled by reads / total reads; this scaled the data so that each locus had equal weight. Actual coordinates were converted to metaplot coordinates based on distance from 5’ basal DCL sites. Scaled depths were normalized by dividing by the number of loci in each set (interface-induced *MIRNA* loci or conserved canonical *MIRNA* loci). Moving averages with a window size of 10 were then plotted (Figure 4E). Transcriptional start sites of interface-induced *MIRNA* primary transcripts were inferred to be the maximum nanoPARE read depth location in the 200 base-pair upstream regions; ties were broken randomly.

### Agroinfiltration of interface-induced *MIRNA* loci

Synthetic double-stranded DNA fragments based on the genomic sequences of two *C. campestris* interface-induced *MIRNA* loci were synthesized: ccm-*MIR12463a* and ccm-*MIR12497f*. Two versions of each were constructed: wild-type, and a version where the ten base-pair USE was randomly scrambled. Synthetic DNA fragments were golden-gate cloned into vector pGGZ001 (Lampropoulos et al., 2013) using sites ‘A’ and ‘G’. The T-DNA regions were confirmed by Sanger sequencing. Complete plasmid sequences are in Supplemental Dataset 5. Plasmids were transformed into *Agrobacterium tumefacians* GV3101-pMP90-pSoup. Agroinfiltration into *Nicotiana benthamiana* leaves was performed as previously described (Liu et al., 2014). Leaf samples from infiltrated areas were collected four days after infiltration and processed for total RNA extraction. Total RNA was extracted using one of the following methods: Tri-Reagent (Sigma), New England Biolabs Monarch Total RNA Miniprep Kit (T2010), or Zymo Quick-RNA Plant Miniprep kit (R2024). RNA samples were diluted to 100ng/ul, and used in stem-loop reverse-transcription reactions (Yang et al., 2014) using primers specific to the target miRNA, and to miR159 (which served as a reference gene). Stem-loop reverse transcription primers were self-annealed at 100uM concentration in IDT Duplex Buffer (100mM potassium acetate, 30mM HEPES pH7.5) by heating in a 94C heat block for two minutes followed by de-activation of the heat block for slow cooling to room temperature. Self-annealed primers were diluted to 50nM stocks in water and stored at −20C. Reverse transcription reactions contained 4ul total RNA (at 100ng/ul), 2ul of a freshly prepared stem-loop primer mixture that had each primer at 5nM concentration, 2ul of 5X reverse transcriptase buffer, 1ul of 100mM DTT, 0.5ul of 10mM dNTP, and 0.5ul of Protoscript II reverse transcriptase (New England Biolabs M0368). Reverse transcription reactions were incubated at 16C for 20 minutes, 42C for 60 minutes, and 80C for five minutes. 1.5ul of the resulting cDNA was used as template in 20ul scale qPCR reactions using Luna Universal qPCR master mix (New England Biolabs M3003) with forward primers specific to the microRNA of interest and a universal reverse primer. PCR conditions for miR12463 detection were 95C for 60 seconds (initial denaturation), 95C 5 seconds (denature), 55C 15 seconds (anneal), 70C 10 seconds (extend and detect). PCR conditions for miR12497 detection were 95C for 60 seconds (initial denaturation), 95C 15 seconds (denature), 52C 15 seconds (anneal), 72C 30 seconds (extend and detect). Reaction efficiencies for each miRNA were calculated by serial dilution of positive cDNA samples. Efficiency-corrected accumulation relative to miR159 was calculated using the method of (Pfaffl, 2001). Primer sequences are in Supplemental Table 4.

## Accession Numbers

Newly created small RNA-seq data have been deposited at NCBI GEO under accessions GSE184641, GSE184642, and GSE205256. Accession numbers for each specific library analyzed are in Supplemental Table 1.

## Supplemental Data Files

**Supplemental Figure 1**. *C. campestris* growth on *A. thaliana* stems.

**Supplemental Figure 2**. *C. campestris* growth on *S. lycopersicum* hypocotyls.

**Supplemental Table 1**. Small RNA-seq datasets used in this study.

**Supplemental Table 2**. *trans*-species microRNA families in *Cuscuta campestris*

**Supplemental Table 3**. Secondary siRNA loci in tomato and *A. thaliana*.

**Supplemental Table 4**. Oligonucleotide sequences

**Supplemental Dataset 1**. Annotations of *Cuscuta campestris* interface-induced *MIRNA* loci. Includes mature microRNAs and stem-loop precursors. Reference genome is assembly version 0.32 from (Vogel et al., 2018), retrieved from https://plabipd.de/project_cuscuta2/start.ep Format: general feature format version 3 (gff3).

**Supplemental Dataset 2**. *C. campestris* interface-induced *MIRNA* hairpin sequences. Format: FASTA.

**Supplemental Dataset 3**. *C. campestris* interface-induced mature miRNA sequences. Both the 5p and 3p miRNAs are included for each locus. These are the actually sequenced miRNA sequences; in a few cases there are polymorphisms between the actual sequences and the corresponding sequence in the reference genome. This likely reflects polymorphisms between the strain of *C. campestris* used in this study relative to the reference genome specimen. Format: FASTA.

**Supplemental Dataset 4**. Annotations of *Cuscuta campestris* conserved canonical *MIRNA* loci. Includes mature microRNAs and stem-loop precursors. Reference genome is assembly version 0.32 from (Vogel et al., 2018), retrieved from https://plabipd.de/project_cuscuta2/start.ep Format: general feature format version 3 (gff3).

**Supplemental Dataset 5**. Sequences and annotations of interface-induced *MIRNA* vectors used in agroinfiltration. Genbank flatfile format (text).

## Acknowledgements

We thank the Huck Institutes Genomics Core Facility for small RNA-seq support and Shawn Burghard for Greenhouse services. This research was supported by awards to MJA from the US National Science Foundation (Award # 2003315) and from the US Department of Agriculture – National Institute of Food and Agriculture (Award # 2018-67013-285).

## Author Contributions

MJA and CH conceived the project and planned the experiments. MJA and CH conducted most experiments, analyzed data, crafted figures, and wrote the manuscript. SM and SG performed small RNA sequencing. JH made the initial discovery of the upstream sequence element.

---

**Supplemental Figure 1.**
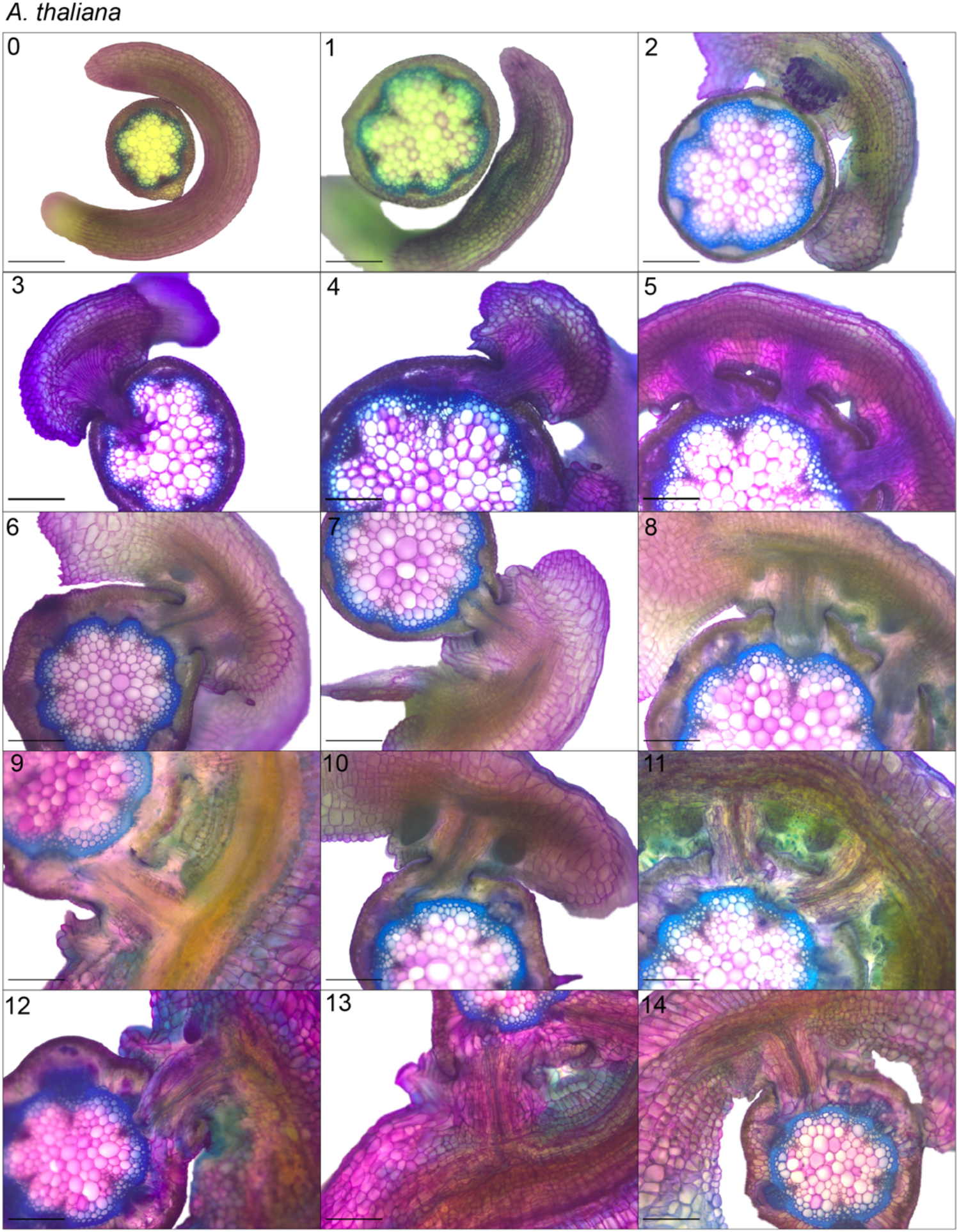
*C. campestris* growth on *A. thaliana* stems. Cross-sections of toludine blue stained vibratome sections for days 0-15 following parasite coiling. Days 2, 3, and 6 were used as Figure 1 A, B, and C, respectively. Scale bars: 200μm.

**Supplemental Figure 2.**
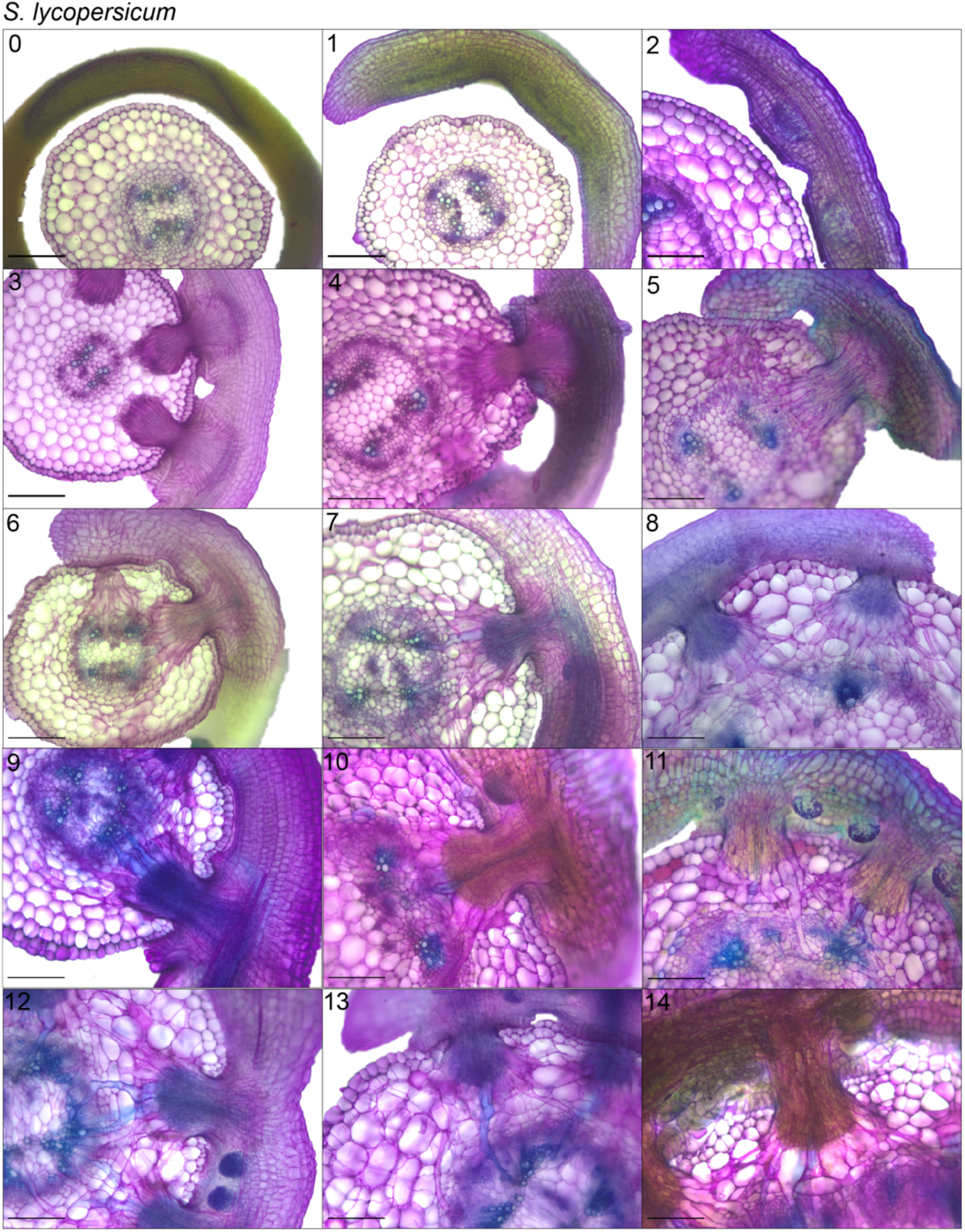
*C. campestris* growth on *S. lycopersicum* hypocotyls. Cross-sections of toludine blue stained vibratome sections for days 0-15 following parasite coiling. Days 2, 3, and 6 were used as Figure 1 D, E, and F, respectively Scale bars: 200μm.

## Footnotes

1 *N.b*. for reviewers and preprint readers: Not all of these annotations are homologs of already described miRBase families, so many don’t yet have miRBase-registered numbers. These cases are temporarily named “ccm-IIM”, for *Cuscuta campestris* interface-induced *MIRNA*s. Upon acceptance, these will be registered at miRBase, and the final miRBase numbers will be substituted into the datasets during final revisions.

